# Unifying Population and Landscape Ecology with Spatial Capture-recapture

**DOI:** 10.1101/103341

**Authors:** J. Andrew Royle, Angela K. Fuller, Christopher Sutherland

## Abstract

Spatial heterogeneity in the environment induces variation in population demographic rates and dispersal patterns, which result in spatio-temporal variation in density and gene flow. Unfortunately, applying theory to learn about the role of spatial structure on populations has been hindered by the lack of mechanistic spatial models and inability to make precise observations of population structure. Spatial capture-recapture (SCR) represents an individual-based analytic framework for overcoming this fundamental obstacle that has limited the utility of ecological theory. SCR methods make explicit use of spatial encounter information on individuals in order to model density and other spatial aspects of animal population structure, and have been widely adopted in the last decade. We review the historical context and emerging developments in SCR models that enable the integration of explicit ecological hypotheses about landscape connectivity, movement, resource selection, and spatial variation in density, directly with individual encounter history data obtained by new technologies (e.g., camera trapping, non-invasive DNA sampling). We describe ways in which SCR methods stand to revolutionize the study of animal population ecology.

## INTRODUCTION

Understanding factors influencing natural variation in population size and structure, demographic rates, and movement has long been a central research focus for population ecologists. Despite well-developed theories over the last half century demonstrating the importance of spatial structure in shaping spatio-temporal population dynamics (e.g., Huffaker 1958, Hanski 1999, Elner et al. 2001), the field of population ecology remains, by and large, unconcerned about within-population spatial processes and their effects on populations. Ecologists routinely study such processes as how individuals use space within their home range, how they perceive connectivity of the landscape, interact with other individuals of the same or other species and how survival or recruitment might be impacted by landscape heterogeneity. However, the population level implications of these processes are not widely studied. Instead the focus is at the individual level, often by studying only a few individuals, with no accounting for how those individuals are sampled from the population. Extending inferences from the individual to the population is not straightforward and in some cases not even possible without a formal statement of a population model linking the sample to the true state, and a description of the sampling process.

Much of what drives the spatial ecological processes that give rise to spatio-temporal population dynamics is the structure and configuration of the landscape (Turner et al. 2001). In fact, linking landscape structure to ecological processes is the primary focus of landscape ecology. This focus on how spatial structure influences ecosystem composition, structure, and function (Turner et al. 2001) by definition, avoids any assumption about spatial homogeneity. When related to animal populations, the tendency in landscape ecology is to focus on movement processes, specifically landscape connectivity (Taylor et al. 1993) and resource selection functions (Chetkiewicz et al. 2009, McLoughlin et al. 2010) rather than demographic rates and population state variables that are of interest in population ecology. Population and landscape ecology offer alternative, yet equally important approaches for understanding spatio-temporal dynamics, yet a consistent theory or quantitative framework linking spatial landscape structure and population ecology does not yet exist.

The ecological theory underpinning landscape and population ecology is well-developed (Tilman & Karieva 1997; Hanski & Ovaskainen 2000, Hanski 2001, Turner et al. 2001, Kot 2001, Williams et al. 2002; Gets & Saltz 2008; Allen & Singh 2016), but testing theoretical models and predictions about spatial ecology in practice is both logistically and statistically challenging. One major impediment is the lack of general mechanistic spatial models that can be applied to empirical data; this precludes rigorous testing of theoretical predictions. Spatial point process models (Illian et al. 2008) provide a natural framework for characterizing the spatial structure of populations assumed to be static and that can be observed with a high degree of accuracy. However, point process models have not been widely adopted in practical field studies of population ecology where individuals cannot be enumerated easily. In practice, populations distributed widely in space must be studied by observing a sample of individuals, sometimes only a very small fraction, at only a few time points and at only a few locations. In some cases, individuals can be continuously monitored (e.g., by telemetry), but in general it is not possible to observe the status of animals perfectly – either their demographic status, their location, or even whether or not they are alive. This is one of the key considerations that has motivated the development and widespread adoption of capture-recapture methods which are now ubiquitous in ecology (Williams et al. 2002; Cooch & White 2006).

For decades, capture-recapture methods have been the cornerstone of ecological statistics as applied to population biology (Nichols 1992, Williams et al. 2002). At their core, capture-recapture models are the canonical class of models for “individual encounter history” data. These data are obtained by capturing or encountering individuals (e.g., using camera traps, acoustic sampling, non-invasive genetic sampling, or direct physical capture), marking them, and observing them over time. Capture-recapture methods have had a profound influence on the study and understanding of demography in wild populations (Karanth et al. 2006, Pradel et al. 2010), in advancing ecological theory (Cooch et al. 2002), and informing modern conservation and wildlife management practices (Nichols and MacKenzie 2004). While capture-recapture has become the standard sampling and analytical framework for the study of population processes (Williams, Nichols & Conroy 2002) it has advanced independent of and remained unconnected to the spatial structure of the population or the landscape within which populations exist. Furthermore, capture-recapture does not invoke any spatially explicit biological processes and thus is distinctly non-spatial, accounting neither for the inherent spatial nature of the sampling nor of the spatial distribution of individual encounters. This precludes the study of many important spatial processes and/or the emerging within-population spatial structure that is arguably as important as demographic rates in population ecology. Recently developed spatial capture-recapture (SCR) methods (Royle et al. 2014) couple a spatio-temporal point process with a spatially explicit observation model which resolves these important criticisms and offers a significant advance in our ability to quantify and study spatial processes using encounter history data. Spatial capture-recapture represents an extension of classical capture-recapture and allows for both the spatial organization of sampling devices and the spatial information that is inherent in essentially all studies of animal populations, i.e., *spatial encounter histories*.

Although a relatively recent advance in the field of statistical ecology (Efford 2004), the past decade has seen an explosive growth in SCR methodological development and applications (Box 1). Spatial capture-recapture provides a quantitative framework that links ecological processes at the individual and population levels. SCR promises the integration of models (hypotheses) of within-population dynamics with "population level" parameters and dynamics. Thus, SCR has proven to be more than simply an extension of a technique, but has emerged as a flexible framework that allows ecologists to test hypotheses about a wide range of ecological theories including landscape and network connectivity (Sutherland et al. 2015, Fuller et al. 2015), demography (Ergon & Gardner 2013, Whittington and Sawaya 2015, Munoz et al, 2016), resource selection (Royle et al. 2013b, Proffitt et al. 2015), and movement and dispersal (Borchers et al. 2014, Lagrange et al. 2014, Schaub & Royle 2015, Royle et al. 2016). While classical capture-recapture methods focus on population level quantities, SCR models allow for the “downscaling” of population structure from coarse summaries (spatial and/or demographic) into finer-scale components by the use of a spatially explicit individual-based point process model. By connecting population level attributes to individual level attributes that are spatially realistic, SCR unifies the fundamental concepts of population and landscape ecology, relating spatial encounters of individuals to explicit descriptions of spatial structure and of how space is sampled (Box 2).

#### Box 1 – New technologies for generating spatial encounter data

The advent of new field-based methodologies for individual identification allows researchers to collect spatial encounter information on individual animals without the need for physically capturing and marking individuals. Additionally, many of the methods are amenable to citizen science approaches (Dickinson et al. 2010) whereby non-professional scientists are engaged in the collection of data (e.g., camera traps, hair snares), providing increased spatial extent of sampling.

*Camera Traps*: With improvements in camera technology, there are many commercially available cameras (a) with superior digital technology that provide still photographs and videos to capture species that are elusive and otherwise difficult to capture. Individuals can be identified from photographs for species that possess distinctive natural marks (e.g., Andean bears (b), tigers (c), wolverines (d), bobcats, jaguars, snow leopards, and others).

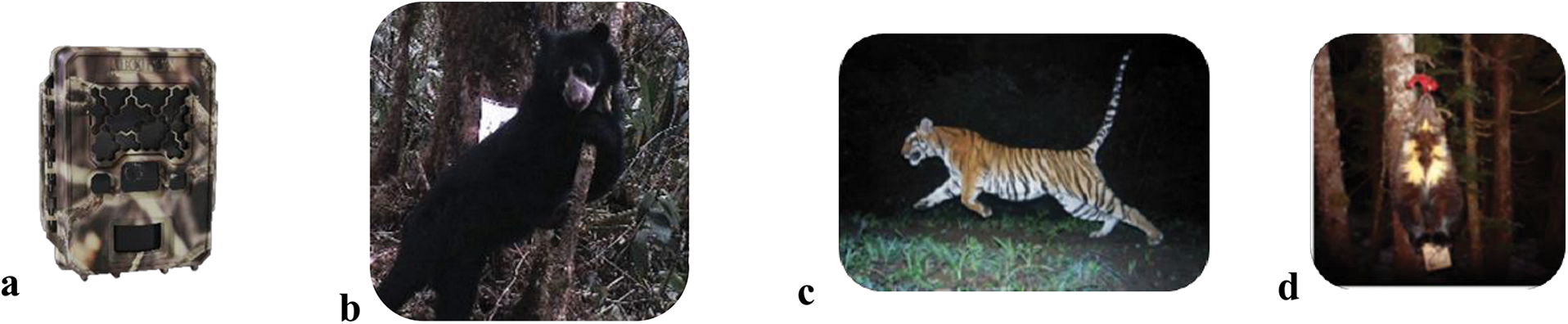

*Non-Invasive Genetic Sampling* (NGS): NGS allows for the identification of individuals without direct observations via the extraction of DNA from samples. Genetic data can be collected from scat, hair, feathers, shed skin, saliva, and urine. Two common methods of obtaining genetic samples are by using devices that snag hair (i.e., hair snares) (a) and scat detection dogs (b). These methods have been employed on marine and terrestrial mammals (e.g., right whales, black bear, fisher, American mink).

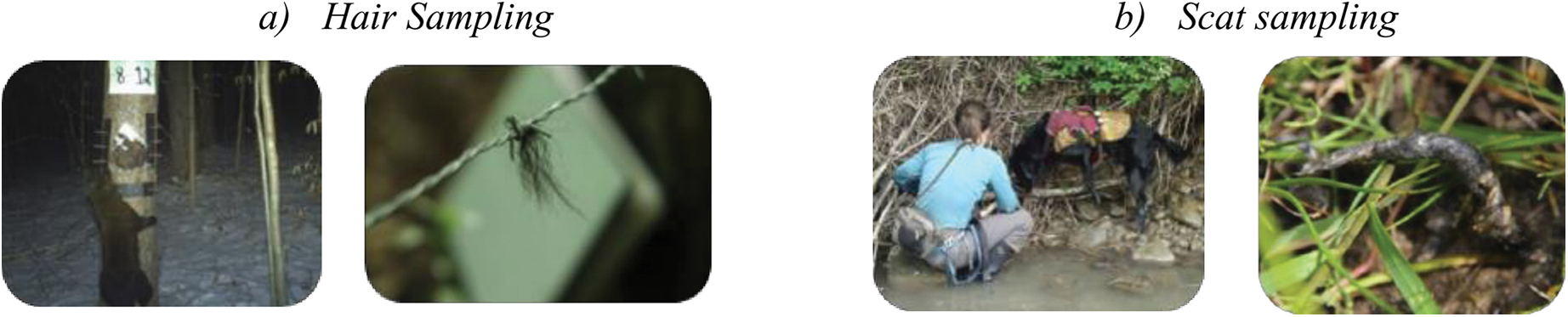

*Bioacoustics*: Spatially separated microphones or hydrophones can be used to detect species that produce sounds for biological purposes such as defending territories, social calling, and mate attracting. Recent advances in bioacoustics technologies and signal detection and recognition algorithms of spectrographs (left) permit the collection of sounds from species such as mammals, birds, and marine mammals.

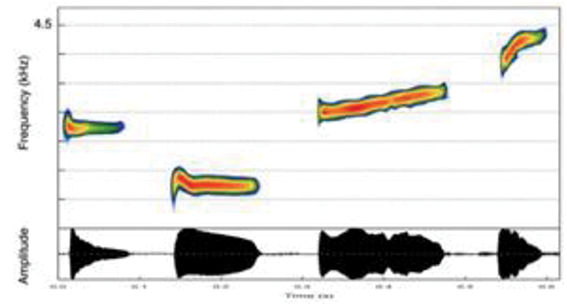

#### Box 2 – Core elements of spatial capture-recapture

*Spatial encounter history data*: Classical capture-recapture summarize spatial data and records only *when* each individual is encountered (a). In practice, data are reduced from a richer 3-d data structure - a record of *when* and *where* each individual was captured (b). Such spatial pattern data are informative about spatial population processes.

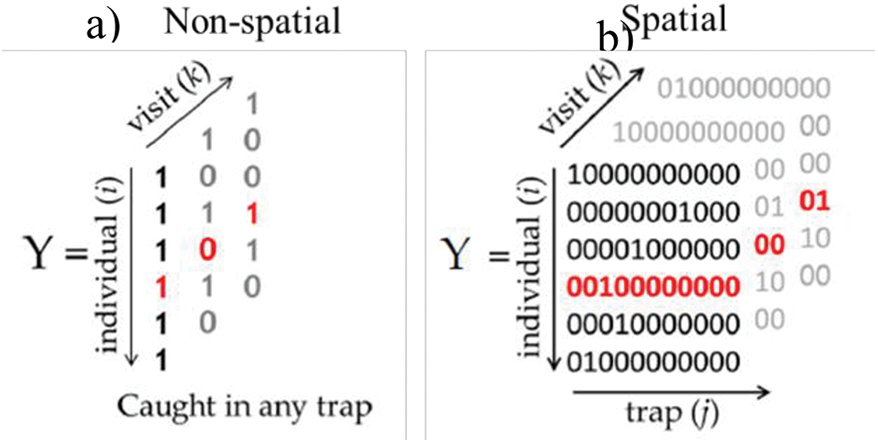

*Encounter probability model*: SCR models describe encounter probability as a function of the distance between a sample location and *s*, the individual’s activity center (the half-normal form is shown to the left). The spatial scale parameter *σ* accommodates individual heterogeneity in detection due to the juxtaposition of individuals with detectors.

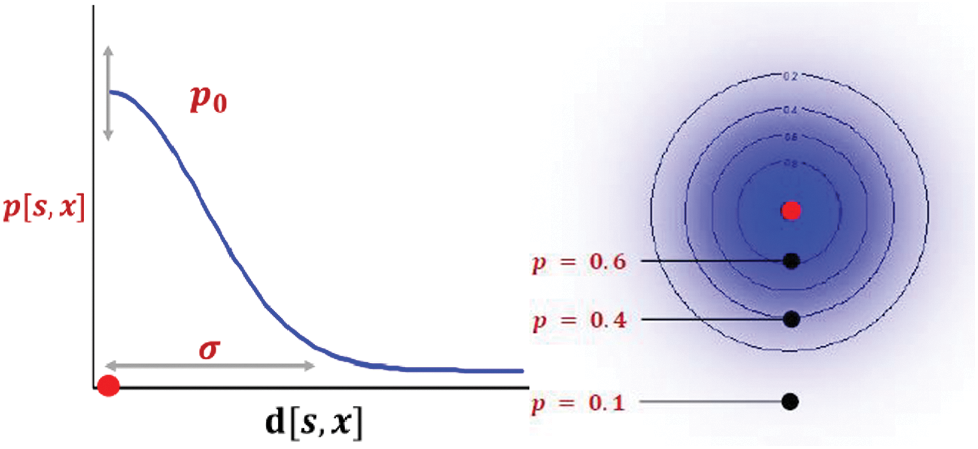

*Spatially explicit point process model*: Encounter histories are modelled conditional on a latent point process describing the spatial distribution of individuals. The null model of uniformity (a) is typically applied and robust to violations. More realistic models allow individuals to be distributed explicitly according to some covariate (b).

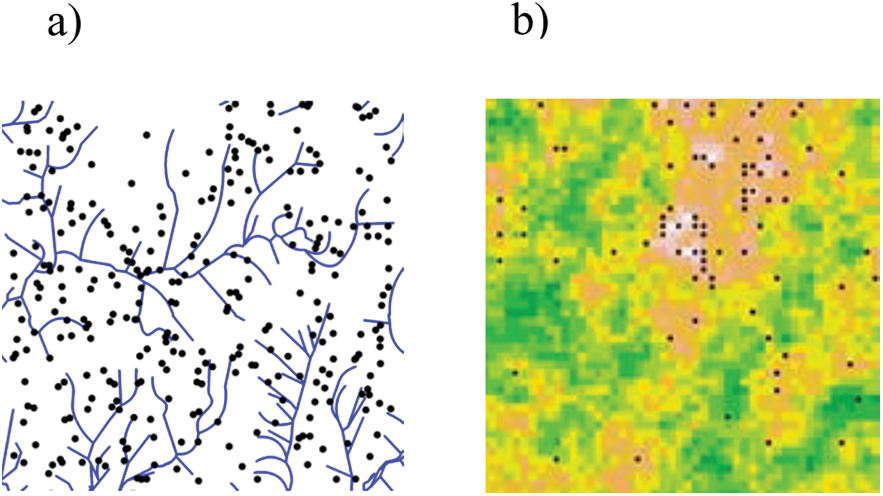

In this review, we describe the basic elements of spatial capture-recapture and how SCR methods advance spatial population ecology by providing a unified framework that integrates important concepts and elements of population ecology and landscape ecology. As such, the framework allows for the study of density, movement, resource selection, landscape connectivity, and other spatial population processes using ordinary encounter history data. Finally, we discuss new directions in the study of animal populations that are made possible by the existence of spatially explicit capture-recapture methods.

## THE ELEMENTS OF SPATIAL CAPTURE-RECAPTURE

Traditional capture-recapture (CR) models were largely motivated by a formal statistical sampling view of how individuals are encountered by sampling, with little or no direct consideration given to the fundamental spatial nature of the sampling. As a result, traditional CR models represent, in essence, “fish bowl" sampling – that is, a system that is devoid of any meaningful spatial context. This leads immediately to several important technical concerns that arise in the application of traditional CR to the study of animal populations which are necessarily spatially explicit.

One important deficiency with classical closed population models is the inability to directly estimate animal *density* (D), arguably the state variable of interest in the vast majority of animal monitoring studies (Krebs 1985, Turchin 1998). This is because, in almost all practical field applications, it is not possible to precisely define the effective area sampled by a set of trapping devices due to movement of individuals into and out of the region within which sampling occurs (Dice 1938; Hayne 1949; Wilson and Anderson 1985a,b). Secondly, the probability of encountering an individual is necessarily heterogeneous among those individuals exposed to sampling. For example, individuals on the periphery of a trapping grid should have lower probability of capture than individuals with home ranges on the interior of the trapping grid (Figure 1). This heterogeneity in encounter probability has long been known to induce negative bias in estimates of abundance (*N*), and hence density (K. Ullas Karanth & Nichols, 1998; Otis, Burnham, White, & Anderson, 1978), and was one of the factors that motivated the development of SCR methods (Efford, 2004). These (and other) technical limitations of the non-spatial CR framework arise directly as a result of a lack of spatial explicitness. On the other hand, SCR integrates models that describe the spatially explicit nature of sampling, how individuals are distributed, and how they use space.

**Figure 1.**
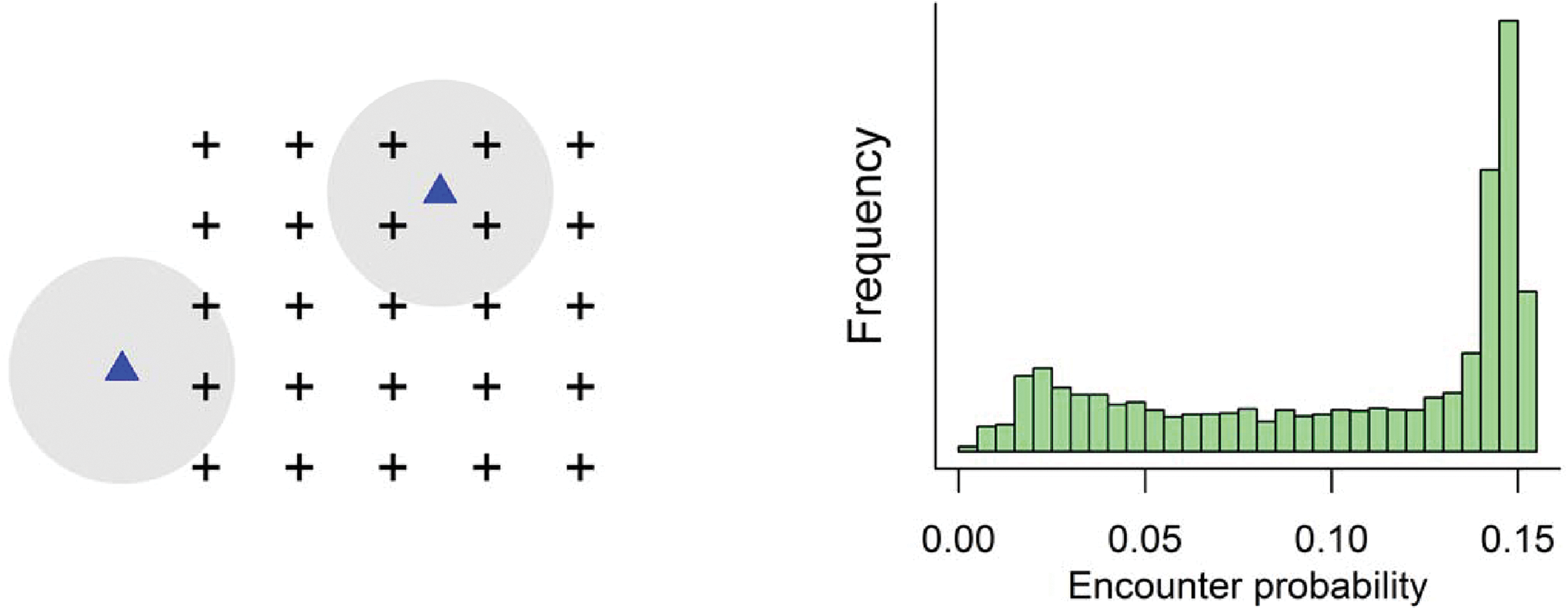
Left: Two home ranges of individuals (gray circles) juxtaposed with a spatial sampling grid (traps) illustrating the variable exposure to trapping based on home range location. Right: the implied distribution of individual encounter probability for a population exposed to sampling by a regular grid (taken from Royle et al. 2014, ch. 5).

SCR models assume that a population of *N* individuals is sampled and that each individual has associated with it a spatial location which represents its activity centre which can be expressed by its *X* and *Y* coordinates as **S***i* = [*S_i,X_, S_i,Y_*]. The collection of activity centres **S**_1_,…,**S***N* can be thought of as the realization of a statistical point process (Illian et al. 2008), a class of probability models for characterizing the spatial pattern and distribution of points. This is perhaps the key innovation of spatial capture-recapture because it is this model that connects observations to much of the ecological theory that can be addressed by SCR. To formalize the point process model it is necessary to describe the probability distribution function of the point locations. The simplest possible point process model is to assume that each of the *N* point locations are distributed uniformly in space (the “uniformity assumption”):

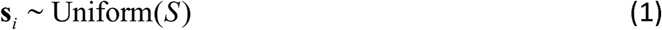

where *S* is an explicit spatial region within which sampling of individuals occurs, and for which inferences about density will be made. Formally this is referred to as the state-space of the point process and is an essential component of a probabilistic characterization of potential activity centres, which are equivalent to individuals in the SCR framework. One important distinction to be made between SCR and classical CR methods is that the state-space *S* is an explicit component of the SCR model. The state-space induces an explicit model of heterogeneous detection probabilities which may affect inferences about density and, hence, population size.

The introduction of this statistical point process – that is, the association of a spatial coordinate with each individual in the population – leads naturally to two distinct conceptually important and powerful modifications of the classical capture-recapture framework which are at the core of the SCR method: First, we can formulate a spatial model for the probability that an individual is captured in each sample location or trap *x_j_* for *j* = 1,2,…,*J*, conditional on its activity centre rather than simply whether an individual was captured at all in a sample occasion, as is the case in traditional CR.

Acknowledging the spatial structure of the traps means observations can be spatially indexed (Box 2, top panel) so encounter histories describe *who* (*i*), *when* (*k*) and importantly *where* (*j*) individuals were encountered, i.e. *y_i,j,k_*. Often, these observations are assumed to be Bernoulli outcomes:

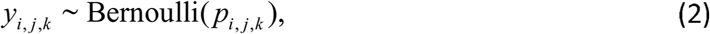

where *p_i,j,k_* is the probability of encountering individual *i* in trap *j* and occasion *k*. It is this model which links the observations (spatially indexed encounters) to the underlying latent point process describing biological pattern and process. At a minimum the encounter probability model depends on distance between the trap location (*x_j_*) and the individual’s activity centre and the individual’s activity centre (*s_i_*) such as the halfnormal encounter model (Box 2, middle panel):

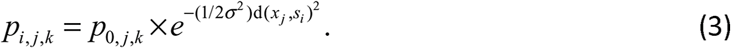

where *p*_0_ is the baseline encounter probability, the probability of encountering an individual at its’ activity centre, the parameter *σ* describes the rate at which detection probability declines as a function of distance, and d(*x_j_,s_i_*) is the Euclidean distance between trap *j* and the activity centre of individual *i* (Box 2, middle panel). In a spatial capture-recapture analysis, the parameters to be estimated are *p*_0_, *σ* and population size *N* or density *D*. We note that the parameter *σ* accommodates individual heterogeneity in detectability but, unlike classical models of heterogeneity (Otis et al. 1978; Dorazio & Royle 2003) the parameter represents an explicit source of heterogeneity, due to the distance between individual activity or home range centres and traps.

The uniformity assumption yields what is usually referred to as a homogenous point process model, although very general models of the point process are possible. For example, when spatially referenced covariates, say *z*(**s**), can be identified that result in spatially heterogeneous density surfaces (Box 2, lower panel), then a standard inhomogeneous point process model posits that

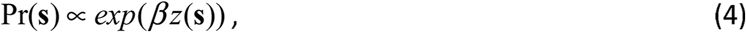

where the parameter *β* corresponds to an explicit hypothesis: “does density depend on the covariate *z*?”

Integrating the point process model with the CR sampling framework leads naturally to a direct focus on inference about parameters of the underlying point process, instead of the abstract quantity N which is devoid of spatial context. Under the SCR modelling framework, the goal is to estimate the number of individuals (or activity centres) within any region of the state-space S. For example, we may estimate density *D*, the number of activity centres per unit area of *S*, or produce predictions of the number of points in any formal subset of *S*, or functions of the entire set of *N* points which might be used to test for spatial randomness, clustering mechanisms (Reich & Gardner, 2014) or other point process assumptions.

For purposes of statistical estimation and inference, the activity centres are regarded as latent variables (i.e., as in classical random effects or mixed models, Laird & Ware 1982). The point process model is then precisely equivalent to the random effects distribution or prior distribution. The resulting model is amenable to analyses by classical methods of statistical inference such as based on marginal likelihood (Borchers & Efford 2008), in which the latent variables are removed from the likelihood by integration, or Bayesian analysis by Markov chain Monte Carlo (MCMC; Royle & Young 2008), in which the activity centres are explicitly estimated along with other unknown parameters and random variables.

SCR models are now routinely applied to many taxa, across a wide variety of systems using a range of sampling methodologies (Box 1). However, the utility of the model reaches far beyond simply estimating density and includes the investigation of important questions about population and landscape ecology which we describe next. Moreover, an enormous number of extensions to SCR models can be accommodated, both to the structure of the ecological processes and the types of observation method, including acoustic sampling (Dawson & Efford, 2009), sampling continuous space instead of using traps (Royle, Kéry, & Guélat, 2011; Royle & Young, 2008), sampling continuous time instead of having discrete sampling intervals (Borchers et al. 2014) and modeling population dynamics such as survival and recruitment using individual level or state-space formulations of classical Jolly-Seber and Cormack-Jolly Seber models (Gimenez et al. 2007; Gardner et al. 2010). We discuss some of these extensions below.

## SCR – A DECADE OF DEVELOPMENT AND APPLICATION

As SCR methods were first appearing more than 10 years ago, the motivation for their development and use was exclusively as a technical device for resolving specific technical limitations of ordinary capture-recapture (Fig. 1). More generally, SCR methods have proved to be a flexible framework for making ecological processes explicit in models of individual encounter history data, and for studying spatial processes such as individual movement, resource selection, space usage, landscape connectivity, population dynamics, spatial distribution, density and inter- and intra-specific interactions. Historically, researchers studied these questions independently, using ostensibly unrelated study designs and statistical procedures.

### SCR for resource selection

SCR models provide a coherent framework for modeling both 2^nd^ and 3^rd^ order resource selection (Johnson 1980; Box 3). Second order resource selection is defined as the processes by which individuals select the location of their home range within a particular landscape. SCR models parameterize an explicit representation of this selection process in the specification of the latent point process model (individual activity centres **s**_1_,…,**s***_N_*). While typical applications involve a relatively simple homogeneous point process model in which activity centers are distributed independently and uniformly over the state-space *S*, the SCR framework accommodates inhomogeneous point process models in which the density of activity centers varies as a function of explicit covariates or flexible spatial response surface models (Borchers & Kidney 2014) that affect density. Inhomogeneous point process models show great promise for testing explicit hypotheses about 2^nd^ order selection, understanding mechanisms that influence species density distribution, and developing conservation and management strategies with explicit abundance- or density-based objectives (Kendall et al. 2015, Proffitt et al. 2015, Sun et al. 2015, Linden et al. 2016).

##### Box 3 – Modeling resource selection with SCR

Resource selection is a multi-scale process (Johnson 1980), determining the range of a species (1st order selection), the distribution of individuals within their range (2nd order), and the use of habitat by an individual within its home range or territory (3rd order). SCR methods allow for explicit modeling of both 2nd and 3rd order resource selection from encounter history data produced from standard capture-recapture methods such as camera trapping, scat and hair sampling for DNA and live trapping.

*2nd order resource selection* is the process that governs the placement or location of individual activity centers. SCR models accommodate explicit models for the probability distribution of activity centers, referred to as inhomogeneous point process models, in which the intensity function depends on landscape or habitat structure:

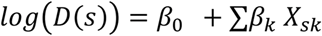

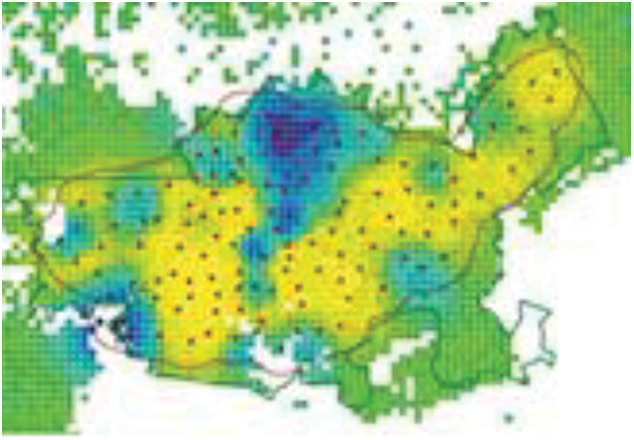

At right, mapped tiger density (from Gopalaswamy et al. 2012).

*3rd order resource selection* affects the SCR encounter probability model (Royle et al. 2013). For some spatially explicit covariate, the probability of encounter can be modeled as a function of both distance and measured covariate with parameter to be estimated:

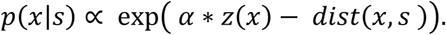

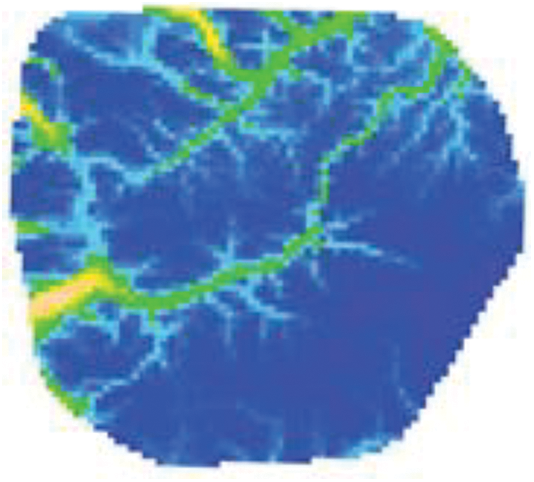

This corresponds to the kernel of standard resource selection models, providing a framework for formal integration of capture-recapture data with data from telemetry studies. At right, estimated encounter probability surface for a black bear population if a trap were placed at a pixel relative to the encounter probability at a pixel of average elevation (from Royle et al. 2013).

Third order resource selection – that is, selection that occurs within an individual’s home range – can be modeled explicitly in SCR models by accommodating habitat structure in the vicinity of sampling (or trap locations) as a covariate that affects the probability of encounter (Royle et al. 2013b). Traditionally, resource selection was studied exclusively by telemetry, and more recently GPS, and because of the high cost are often based on a small sample of individuals observed many times. Conversely, SCR methods may produce a sample of many more individuals, and direct information about population level resource selection from spatial encounter data. However, individuals in the population often do not need to be physically captured to obtain this information by SCR methods (e.g., by camera trapping). Thus, SCR provides an alternative, efficient, and cost effective framework for studying the important population process of resource selection from individual encounter history data for species that classical telemetry may not be viable and that is based on a larger sample of the population.

### SCR for modeling movement and dispersal

The direct linkage of the SCR encounter probability model to movement of an individual within its home range is one of the basic concepts of SCR (Royle & Young 2008; Borchers et al. 2014). However, one of the key assumptions of most SCR methods to date is that the latent point locations which represent the individual activity centers are static variables. In a sense this is a manifestation of type of population “closure”; individuals are allowed to move around in space, but their expected location is assumed not to change over the course of the study. Recent attention has been given to modifying the underlying state point process model to accommodate temporal dynamics such as dispersal or transience (Schaub & Royle 2014; Ergon & Gardner 2014). These models formally allow for the estimation of survival probability that is free of biasing effects of dispersal whereas, classically, only “apparent survival” has been estimated from standard capture recapture data (Schaub et al. 2004). Even in populations where mortality or recruitment are absent, including a dynamic spatial process to account for dispersal and transience is possible by coupling a latent movement model with a spatial model of the encounter process (Royle et al. 2016). For example, to modify the point process model to allow for an individual’s activity center to shift from time *t* – 1 to time *t* we might accommodate this with a simple Markovian movement model where the difference between successive activity centers has variance *τ*^2^:

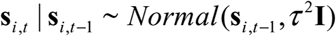

Thus, SCR has a characterization as a state-space (Patterson et al. 2008) or hidden Markov model (HMM; Langrock et al. 2012) with specific forms of observation model governed by spatial sampling and an underlying latent process model that describes movement of individuals on the landscape. As a result, SCR offers a general framework for the formal study of dispersal, transience, and other types of movement from individual encounter history data.

### Modeling landscape connectivity

One of the core elements of SCR is the model for encounter probability which we described above as a function of Euclidean distance between activity centers and sample locations (Box 2). However, the Euclidean distance assumption implies a simplistic model of space usage – that individual home ranges are symmetric and stationary. In practice, we expect individual home ranges to be influenced by local landscape characteristics and structure. One approach for accommodating this landscape structure-induced asymmetry in space use in SCR models is the relaxation of the Euclidean distance assumption.

This is achieved using an alternative distance metric that is related to the landscape through which distance is being measured, thus allowing the degree of asymmetry to be estimated using a model that relates the observed spatial pattern of observations explicitly to the measurable landscape characteristics. For example, Royle et al. (2013a) suggested using least-cost path distance with the exception that, rather than being defined *a priori* based on opinion as is customary (Zeller et al. 2012), the resistance parameters are estimated using standard likelihood methods based on spatial encounter histories (Box 4). This idea was extended by Sutherland et al. (2015) for highly structured dendritic networks, an extreme yet intuitive conceptual setting for investigating the utility of this asymmetric space use model. The major development is that the model of asymmetric space use can be used to jointly estimate density and resistance parameters which are typically defined a priori based on opinion (Zeller, McGarigal & Whitely 2012), yielding ecologically interesting and realistic individual home range geometries which can be scaled up to the landscape level based on direct estimates of the landscape structure-space use relationship (Box 4). What results is the important notion that ordinary encounter history data that is extensively collected in ecological studies with relative ease can now be used to formally characterize landscape connectivity within a framework of statistical inference. Sutherland et al. (2015) defined several intuitive measures of landscape connectivity based on such upscaling based on the SCR encounter probability model to the landscape scale. The asymmetric space use model described here and in Box 4 in general, can be extended to multiple landscape characteristics as would be done with any log-linear regression model, and requires only that the landscape covariates are defined at the pixel level. While current applications have focused on river networks (Fuller et al. 2016), the approach should be highly relevant for any species for which one or more landscape features act to impede or facilitate movement (Morin et al. in press) (e.g., extreme topographic variation, well-defined networks of roads and trails used by a species, Box 4). Moreover, SCR offers a formal model-based solution for investigating the strength of landscape interactions, avoiding the need to arbitrarily prescribe resistance values. The possibility exists to consider other non-Euclidean distance metrics such as circuit distance (McRae et al. 2006) or flexible deformations of geographic space (Sampson & Guttorp 1992).

##### Box 4: Esiimating landscape Connectivity

*Landscape structure* influences local movements such that, during monitoring, the pattem of individual observations around its activity center is likely to deviate from the assumption of a circular home range. Understanding patterns of space use, and thus estimating encounter rates without bias, reqiures that the structure of ecological landscapes is explicitly accounted for. Using a least cost path approach, SCR allows the estimation of one or more resistance Parameters, *δ*, characterizing how movement is influenced by landscape structure. Euclidean distance in Eqn. 3 and Box 2 is replaced by *ecological* distance, *d*_ecol_, the length of the least cost path between two points (*v*_0_ and *v_T_*):

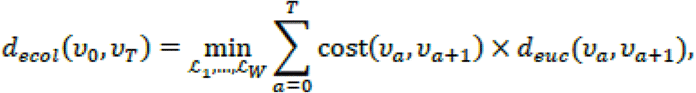

where *L_w_* (*v*_0_, *v_T_*) = {*v*_0_,…,*v_T_*} denotes any path consisting of *T* adjacent lines connecting adjacent pixels, and cost(*v_a_*, *v*_*a*+1_) is a cost function which is a log linear function of the average pixel-specific covariate values:

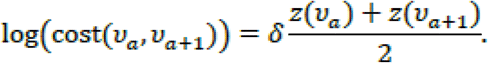

Estimation of the resistance parameter provides a direct measure of the strength of species…landscape interactions which has important implications. First, this is a model for asymmetric space use that simultaneously relaxes assumptions of symmetry and stationarity, allowing *home range geometry* to vary dependnig on location and local landscape structure. Secondly, for a known landscape, the probability of use for any pixel on the landscape can be computed given an individual’s location, i.e., individual local Connectivity, and it follows therefore that the composite local connectivity surfaces for any collection of activity centers provides a *model-hased landscape connectivity* surface informed by the estimate of the species-landscape parameter.

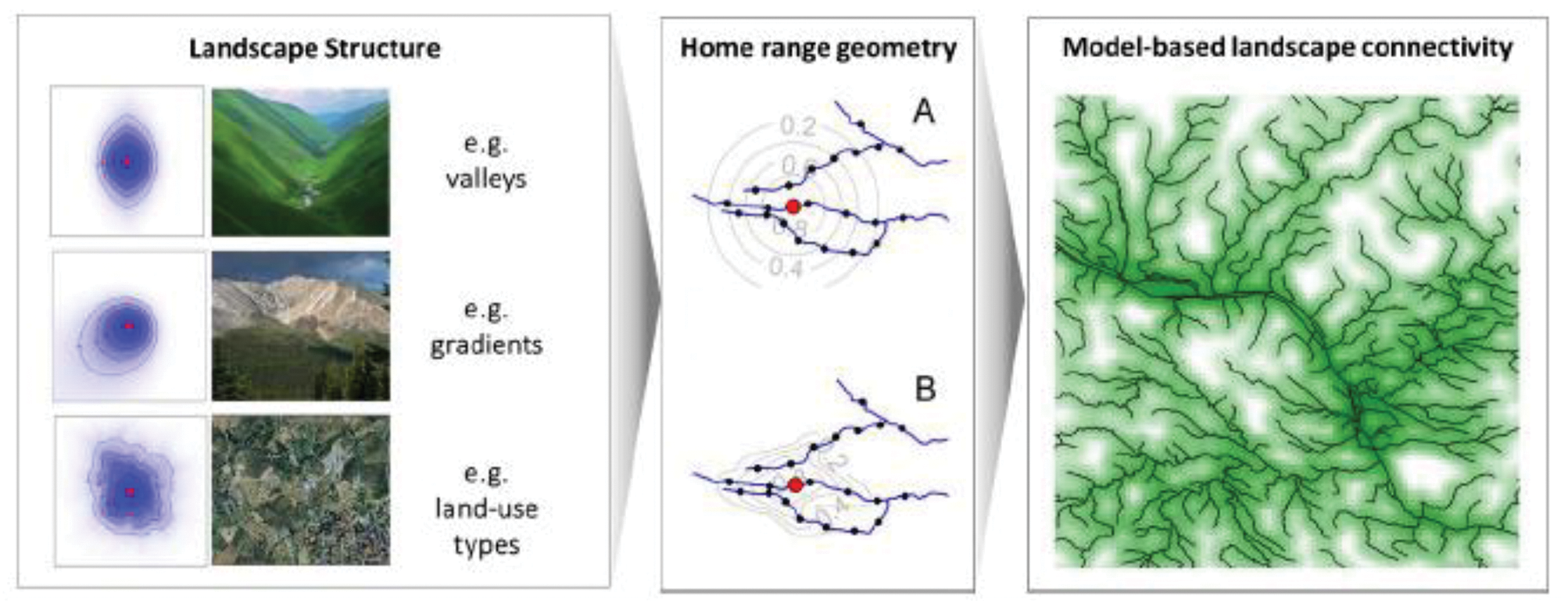

## FUTURE DIRECTIONS OF SPATIAL CAPTURE-RECAPTURE

The relevance of SCR methods is expanding rapidly because these techniques allow ecologists to explicitly test hypotheses about the mechanisms that drive ecological phenomena as diverse as habitat selection, persistence of rare species, community assembly, invasion, and genetic diversity. The developments described above represent significant contributions to applied population ecology despite their relative infancy, and we believe the potential for SCR in ecology has not yet been fully realized. We highlight specific and potentially fruitful development areas for SCR that have the potential to make further contributions with regard to wildlife population sampling, and/or developing and testing ecological hypotheses.

### Landscape management and corridor design

It is possible to use SCR with individual encounter history data to inform landscape management decisions such as related to stepping stone and corridor design problems because SCR models provide spatially explicit within-population information about density and abundance, thus providing objective inferences about where the population is distributed in space and why. When combined with explicit models of connectivity (previous section), spatially explicit metrics which integrate information about both density and connectivity (Sutherland et al. 2015; Fuller et al. 2016; Morin et al. 2016) can be estimated, thus providing information about quality of the landscape for maintaining connectivity and also for maintaining source populations of important species.

Corridors are increasingly used as conservation tools, designed to facilitate movement of individuals between habitat patches, or between two nodes or habitat blocks separated by some distance (e.g., two protected areas) with the ultimate goal of maintaining landscape connectivity. In the most general sense, corridor design involves defining a resistance value (i.e., resistance of the landscape to animal movement) of each pixel in the landscape as a function of pixel characteristics, and then subsequently selecting the lowest cost pixels, typically evaluated by estimating the cumulative cost of moving from one area to another. The resistance of a landscape is approximated by a ‘cost’ value, representing how difficult it is for an individual to move through a landscape. High quality habits are more permeable to movement and infer lower ecological costs (i.e., they provide increased survival and reproduction) relative to lower quality habitat. Resistance values are most often based on subjective expert opinion or data from previously published studies (Zeller et al. 2012). These user defined resistance models have been tested based on limited inference from few radiomarked individuals (Driezen et al. 2007). However, examples exist of deriving resistance values from occurrence probability from occupancy models (Walpole et al. 2012) or using a variety of different threshold values based on the most traversable habitat from radio-marked individuals (Poor et al. 2012). Of importance is that these applications fail to utilize the information from animal movements to directly estimate landscape resistance values.

We are aware of only one application of using capture-recapture data for formal inference about landscape resistance for a species (Fuller et al. 2016), which was based on the Ecological distance SCR model (Royle et al. 2013a, Sutherland et al. 2015). Further, corridor conservation has been devoid of explicit consideration of local population density. SCR models allow for the simultaneous estimation of the two processes that are most critical to the conservation of spatially-structured populations, density and connectivity. Morin et al. (in press) derived a model-based estimator of landscape connectivity (i.e., density-weighted connectivity) that estimates both the spatial distribution and connectivity of individuals across a landscape. Spatial optimization approaches that maximize density-weighted connectivity would identify areas on the landscape that support the highest number of individuals and best landscape connectivity and would therefore have the greatest potential for application in corridor conservation and landscape management.

### Modeling spatial interactions

The latent point process describing the spatial distribution of individuals is a central component of SCR methods. Parameterization of this point process allows encounter history data to be used to develop models that explicitly address theories related to competition, including territoriality (Reich & Gardner 2014) and maintenance of coexisting species and species diversity. Extending the point process model to account for dependencies among multiple species simultaneously occupying a landscape may provide an analytic framework for the empirical study of inter and intra-specific competition and landscape level spatial structure in species assemblages. This should have enormous relevance in understanding host-pathogen and disease in natural systems where transmission depends on local interactions of individuals and local density. Where individuals live and who they interact with are fundamental elements contributing to the dynamics of disease and pathogen systems.

### Acoustic sampling

Acoustic sampling is emerging as a promising technology for sampling vocal species such as birds, anurans, marine mammals, and primates, and the application of these methods is increasing rapidly (Marques et al. 2009; Blumstein et al. 2011). Information on signal strength and/or direction gives imperfect information about the source of the vocalization although statistically pinpointing the source has been recognized as being analogous to inference about the activity center in SCR methods, and therefore SCR has been adapted to accommodate data obtained by acoustic sampling methods (Dawson & Efford 2009; Efford et al. 2009; Borchers et al. 2015; Stevenson et al. 2015; Kidney et al. 2016). It seems probable that these technologies will become the *de facto* sampling method in bird population studies due to the increasing affordability of the technology.

### Uncertain identity

Given the widespread adoption of non-invasive sampling technologies, which may only yield partial information on the identity of individual samples, it will become important to accommodate uncertainty in individual identity in to studies of animal populations that use individual encounter histories. There has been considerable attention paid to the problem of uncertain identity in capture-recapture (Link et al. (2010), Bonner & Holmberg (2013), McClintock et al. (2013)). However, such methods have developed in the context of classical capture-recapture methods which ignore the spatial information inherent in most animal population sampling studies. On the other hand, for most populations we should expect that the spatial location of samples should be informative about the uncertain identity of those samples (Chandler & Royle 2013, Chandler & Clark 2014, Royle 2015, Augustine et al. 2016). That is, all other things being equal, spatial samples that are in close spatial proximity to one another should more likely be of the same individual than samples that are far apart. Thus, dealing effectively with an uncertain identity of an individual is fundamentally a spatial problem for which SCR offers a solution.

Methods of accommodating uncertain identity and partially marked populations are promising avenues for the formal integration of citizen science data collection with population ecology studies based on capture-recapture. SCR facilitates the use of citizen science in studies of population ecology because citizen science schemes naturally produce abundant information about individual location, potentially useful in spatial mark-resight and similar SCR models. Thus, involving citizens in data collection will produce large quantities of confirmations of species and their locations, but potentially no individual identity of the observations.

## CONCLUSIONS

Two technological advances have influenced the present and future of animal population ecology in a way that we believe is more profound than any advance in quantitative ecology since the invention of computers. First is the development of new technologies for obtaining spatial encounter information on individuals (Box 1). These technologies have revolutionized applied population ecology. Simultaneous to the development of these new field techniques has been the increasing spatialization of ecological process models seen in the advancing fields of landscape ecology and metapopulation ecology, along with the increasing utilization of statistical point process models throughout population ecology. SCR lies at the convergence of these two technological advances, combining a spatially explicit observation model that describes data collected using new technologies such as noninvasive genetics or camera trapping, with spatially explicit models describing how individuals are distributed across a landscape.

Estimating abundance or population size is one of the most important problems in applied ecology permitting the evaluation of sophisticated questions in population dynamics (Krebs 1984, Williams et al. 2002, Sutherland et al. 2013) and providing necessary information for the conservation and management of important species (Karanth & Nichols, 2002). SCR has become the standard method for obtaining such information for many species, and is now routinely used to estimate abundance of populations of conservation concern including species such as tigers (Royle et al. 2009), grizzly bears (Efford & Mowat 2014; Kendall et al. 2015), wolverines (Magoun et al. 2011, see Box 1) and jaguars (Sollmann et al. 2011). These and many other species are extremely difficult to capture and so non-invasive sampling combined with SCR methods are perfectly suited to study these species. Moreover, many populations exist in such low densities that obtaining sufficient sample sizes of individuals can be challenging, and thus making the most efficient use of all data, and in particular, spatial recaptures which are discarded by ordinary capture-recapture, is critically important.

While SCR methods were developed originally as a tool for inference about animal density from capture-recapture data (Efford, 2004) motivated by the need to address specific technical limitations of ordinary capture recapture methods (Fig. 1), they have proven to be more than simply an extension of technique. Spatial capture recapture has profoundly affected the manner in which capture-recapture is used in studies of animal populations because they allow testing of explicit hypotheses of core elements of population and landscape ecology by formally integrating technical descriptions of these processes with encounter history data obtained by sampling animal populations. SCR models include an explicit model of density and thus relationships can be modeled between density and landscape features or other population attributes. For example, SCR models can be used to test hypotheses related to density dependence in animal populations, such as the relationship between density and home range size (Efford et al. 2015). In addition, understanding movement of individuals over the landscape is a key objective throughout ecology, and SCR enables the formal integration of explicit movement, dispersal, and survival models with models of density and other population characteristics (Ergon & Gardner 2015; Schaub & Royle 2015). Because SCR models are spatially explicit, we believe it is possible to consider explicit modeling of population dynamic rates as a function of local density. Finally, landscape connectivity is a fundamental element of landscape ecology and explicit models of connectivity can be integrated directly with models of spatial encounter history data within the SCR framework to provide population-level estimates of connectivity parameters. Sutherland et al. (2015) and Fuller et al. (2016) develop SCR models in highly structured landscapes and demonstrate formal inference for an explicit model of landscape connectivity and resistance, estimated from individual encounter history data from a capture-recapture study of mink.

SCR is not only revolutionizing how population and landscape ecology questions can be addressed but also in the way we observe populations. For example, use of scat dogs to sample space using unstructured area searches is extremely practical for studying many species and this method has grown rapidly in recent years. When data are obtained in this manner, it is imperative that the spatial structure of sampling be accounted for and SCR accommodates this by explicitly using GPS search lines in place of trap locations. Acoustic sampling (Dawson & Efford 2009) is a promising new technology for sampling birds and many other species. However, without a spatially explicit model that describes both the sampling and underlying process, it is not possible to connect observed acoustic encounter data to meaningful biological parameters such as population density. Finally, SCR has potential as the framework for integrating individual encounter data with inexpensive, broad scale auxiliary data such as from occupancy studies (Chandler & Clark 2014; Whittingham & Chandler in press) and potentially even citizen science programs.

At the core of science is the notion of testing theories by confronting models with data. At the level of the population this has been recognized as a promise of capture-recapture for several decades (Nichols 1992), but the use of capture-recapture has not been widely used to address questions related to within-population spatial structure and population dynamics. However, SCR achieves this promise, by integrating a formal spatial model describing how individuals are distributed over a landscape, with a formal spatial model for how the population is sampled. SCR enables testing explicit spatial mechanisms and processes and improves understanding of spatial ecology from individual encounter history data. SCR incorporates elements of population structure and dynamics and explicit spatial and landscape structure to provide a quantitative framework that unifies population and landscape ecology.

